# Detecting time-varying genetic effects in Alzheimer’s disease using a longitudinal GWAS model

**DOI:** 10.1101/2023.10.17.562756

**Authors:** Xiaowei Zhuang, Gang Xu, Amei Amei, Dietmar Cordes, Zuoheng Wang, Edwin C. Oh, Alzheimer’s Disease Neuroimaging Initiative

## Abstract

**Background:** The development and progression of Alzheimer’s disease (AD) is a complex process that can change over time, during which genetic influences on phenotypes may also fluctuate. Incorporating longitudinal phenotypes in genome wide association studies (GWAS) could help unmask genetic loci with time-varying effects. In this study, we incorporated a varying coefficient test in a longitudinal GWAS model to identify single nucleotide polymorphisms (SNPs) that may have time- or age-dependent effects in AD.

**Methods:** Genotype data from 1,877 participants in the Alzheimer’s Neuroimaging Data Initiative (ADNI) were imputed using the Haplotype Reference Consortium (HRC) panel, resulting in 9,573,130 SNPs. Subjects’ longitudinal impairment status at each visit was considered as a binary and clinical phenotype. Participants’ composite standardized uptake value ratio (SUVR) derived from each longitudinal amyloid PET scan was considered as a continuous and biological phenotype. The retrospective varying coefficient mixed model association test (RVMMAT) was used in longitudinal GWAS to detect time-varying genetic effects on the impairment status and SUVR measures. Post-hoc analyses were performed on genome-wide significant SNPs, including 1) pathway analyses; 2) age-stratified genotypic comparisons and regression analyses; and 3) replication analyses using data from the National Alzheimer’s Coordinating Center (NACC).

**Results:** Our model identified 244 genome-wide significant SNPs that revealed time-varying genetic effects on the clinical impairment status in AD; among which, 12 SNPs on chromosome 19 were successfully replicated using data from NACC. Post-hoc age-stratified analyses indicated that for most of these 244 SNPs, the maximum genotypic effect on impairment status occurred between 70 to 80 years old, and then declined with age. Our model further identified 73 genome-wide significant SNPs associated with the temporal variation of amyloid accumulation. For these SNPs, an increasing genotypic effect on PET-SUVR was observed as participants’ age increased. Functional pathway analyses on significant SNPs for both phenotypes highlighted the involvement and disruption of immune responses- and neuroinflammation-related pathways in AD.

**Conclusion:** We demonstrate that longitudinal GWAS models with time-varying coefficients can boost the statistical power in AD-GWAS. In addition, our analyses uncovered potential time-varying genetic variants on repeated measurements of clinical and biological phenotypes in AD.

## 1. Background

Alzheimer’s disease (AD) is the most common form of age-related dementia. The biological constructs and hallmark pathologies in AD are characterized by extracellular β-amyloid protein deposition, intraneuronal pathological tau protein accumulation, accompanied by neurodegeneration and neuroinflammation[1, 2]. As a highly heritable disorder, genetic factors contribute significantly to the development of AD. The heritability of AD is approximately 60% to 80%, first estimated from genetic twin studies[3] and later corroborated with large-scale genome-wide association studies (GWAS)[4, 5]. Delineating the strong genetic component in AD has become a major objective in AD research, as it provides an opportunity to 1) understand the disease etiology and risks, 2) characterize pathophysiological pathways, and 3) identify biological features as potential diagnostic and prognostic markers.

To date, large-scale GWAS have reported more than 80 putative AD-associated loci and genes[4–8], among which, the *Apolipoprotein E (APOE)* E4 allele on chromosome 19 has been shown to have the largest genetic risk towards AD. Conventionally, AD GWAS have focused on single time point measurements with less emphasis on dynamic phenotypes in a repeated measures setting. Interestingly, longitudinal studies have shown that the genetic architecture of gene expression regulation can be unstable over time and linked with aging[9]; in addition, the genetic contributions towards complex traits such as BMI[10] and hypertension[11] also exhibit time- or age-dependent variability. Pertinent to AD, studies on *APOE* have shown that E4 allele counts demonstrate an inconstant hazard in developing AD, which declines with increasing age[12]. Therefore, as an aging and complex disorder, AD may have critical timelines for the onset and progression, during which genetic influences on phenotypes may also fluctuate. Thus, our main objective of this study was to identify time-varying genetic contributions to phenotypes in AD, with the expectation that the identified SNPs could further assist in delineating genetic mechanisms relevant to AD.

Besides focusing on single time point measures, current AD GWAS have also been conducted using clinically diagnosed case-control subjects. Following the amyloid-tau-neurodegeneration (ATN) framework, about 20%-40% clinical AD cases do not have amyloid pathology and are non-biological AD cases[1]; therefore, non-AD pathology might significantly contribute to AD genetic signals in traditional GWAS. Additionally, association studies with other ATN biomarkers, or within biologically-defined AD participants, might be restricted by the limited sample size and thus suffer from reduced statistical power. Longitudinal models in this case could take advantage of repeated phenotypic measures from the same subject to potentially boost the statistical power in association analyses, which could in turn allow GWAS to be performed on ATN-specific biomarkers, or within biologically-defined AD participants. Therefore, in the current study, we focused on both clinical and biological phenotypes using longitudinal GWAS models.

More recently, several large-scale data initiatives in AD have collected and measured longitudinal diagnoses and ATN biomarkers over time[13–15]. Alzheimer’s neuroimaging data initiative (ADNI, http://www.adni-info.org) is one of these databases, which is a multicenter, multi-phase study assessing clinical, imaging, and genetic biomarkers in AD. Such data enable the community to investigate time-varying genetic effects on various clinical and biological phenotypes over the complex disease course in AD.

Several statistical models have been applied in longitudinal GWAS. Among which, linear mixed effects models and empirical Bayes models are most commonly used to account for dependence structure in longitudinal observations from the same participant[16]. Using these methods, studies have reported novel variants and genes associated with disease progression or cognitive resilience in AD[17, 18]. Varying coefficient models, as an extension of generalized mixed effects models, have been designed to specifically capture the time-varying genetic effects on dynamic traits in longitudinal GWAS[11]. These varying-coefficient models in longitudinal GWAS have led to the identification of time-dependent genetic effects in cocaine users[19], subjects with hypertension[11], and hippocampal volumes in AD subjects[20].

In this study, we applied a varying coefficient model to perform longitudinal GWAS on both a binary phenotype of clinical impairment status and a continuous phenotype of brain amyloid accumulation in AD. We hypothesized that, with increased statistical power and modeling of varying coefficients, longitudinal GWAS models can support the detection of time-varying genetic effects in repeatedly measured phenotypes. We expect our results to improve the identification of genetic variants associated with fluctuating pathological or clinical phenotypes in AD.

## 2. Methods

We performed longitudinal GWAS using a retrospective varying coefficient mixed model association test (RVMMAT), and Fig. 1 depicts our overall pipeline. Data from ADNI and the National Alzheimer’s Coordinating Center (NACC) were utilized as the main and replication datasets, respectively. Both ADNI (http://www.adni-info.org) and NACC (https://naccdata.org/) are publicly available databases that can be accessed upon reasonable requests on their websites. No approval or participant consent was obtained locally from these participants.

**Fig. 1.**
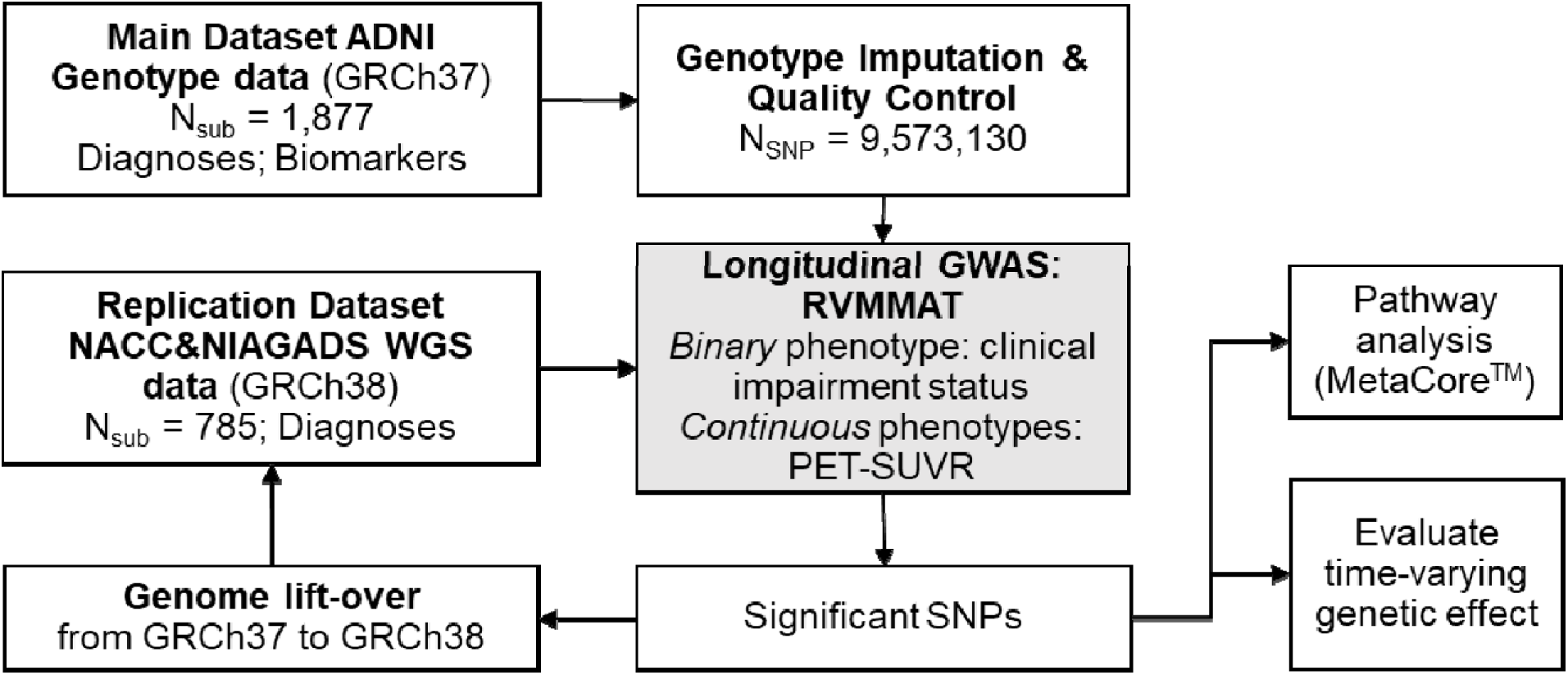
Analyses pipeline. RVMMAT was first performed to identify significant SNPs for both binary and continuous phenotypes using ADNI participants (center grey boxes). Post-hoc analyses on significant SNPs were then performed, including identifying 1) enriched functional pathways and 2) time-varying genetic effect on phenotypes. Replication analyses were next performed on significant SNPs using NACC participants.

### 2.1 Primary dataset: ADNI participants

#### Genotype data

We included a total of 1,877 older participants with genome-wide genotyping data available from the ADNI database. Participants’ demographics including sex and race were downloaded from the ADNI website and listed in Table 1. Among 1,877 participants, 1,020 were men and 857 were women; 74 were African Americans (3.94%), 1,745 were European Americans (92.97%), and 58 were of other races (3.09%). In addition, the 1,877 participants were genotyped in ADNI1, ADNIGO/2 and ADNI3 phases separately. ADNI1 samples (N_sub_=757) were genotyped using the Illumina Human610-Quad BeadChip, with 620,901 genome-wide single nucleotide polymorphisms (SNPs). ADNIGO/2 samples (N_sub_=793) were genotyped using the Illumina HumanOmniExpress BeadChip, with 730,525 genome-wide SNPs. ADNI3 samples (N_sub_=327) were genotyped using the Illumina Infinium Global Screening Array v2, with 759,993 genome-wide SNPs. Genotyping data in PLINK format were downloaded from the ADNI website, and all genotype files were aligned to genome reference consortium human build 37 (GRCh37)[21].

**Table 1.**
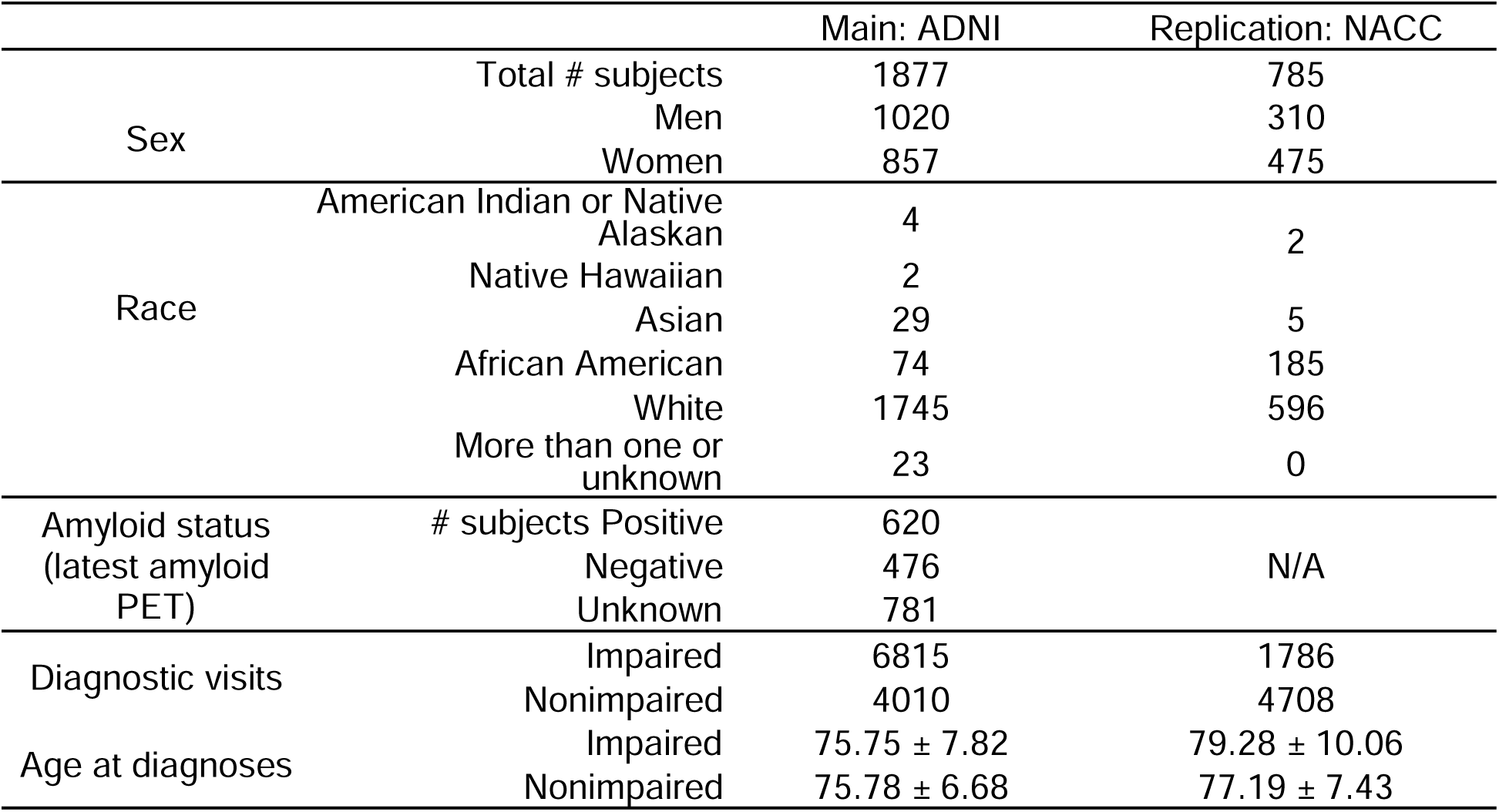
Demographics of ADNI (main dataset) and NACC (replication dataset) participants.

|  |  | Main: ADNI | Replication: NACC |
| --- | --- | --- | --- |
| Sex | Total # subjects | 1877 | 785 |
|  | Men | 1020 | 310 |
|  | Women | 857 | 475 |
| Race | American Indian or Native Alaskan | 4 | 2 |
|  | Native Hawaiian | 2 |  |
|  | Asian | 29 | 5 |
|  | African American | 74 | 185 |
|  | White | 1745 | 596 |
|  | More than one or unknown | 23 | 0 |
| Amyloid status<br>(latest amyloid PET) | # subjects Positive | 620 |  |
|  | Negative | 476 | N/A |
|  | Unknown | 781 |  |
| Diagnostic visits | Impaired | 6815 | 1786 |
|  | Nonimpaired | 4010 | 4708 |
| Age at diagnoses | Impaired | 75.75 ± 7.82 | 79.28 ± 10.06 |
|  | Nonimpaired | 75.78 ± 6.68 | 77.19 ± 7.43 |

#### Genotype data imputation and quality control

Genotyping data were organized and prepared using the imputation preparation and checking tool from the McCarthy Group (https://www.well.ox.ac.uk/~wrayner/tools/). The pre-processed genotyping files were uploaded to the Michigan imputation server[22] and imputed using Mimima4, with the 1000 Genomes phase 3 data (European) as a reference panel. We next tested Hardy-Weinberg equilibrium at each SNP, and SNPs meeting the following quality-control conditions were retained: (1) call rate > 99%, (2) Hardy-Weinberg χ^2^ statistic *p-value* >10^-6^, and (3) minor allele frequencies > 1%. Taken together, a final set of 9,573,130 SNPs were examined in the following longitudinal GWAS analyses.

#### Longitudinal binary phenotype: clinical impairment status

For each participant, we obtained their clinical diagnoses at each visit from all ADNI phases (DXSUM_PDXCONV_ADNIALL.csv file from the ADNI website). There was a diagnostic-criteria change from ADNI1 to ADNIGO/2 phases, and therefore, clinical visits with reverse diagnoses from ADNI1 to ADNIGO/2 were not included in the following analyses.

Collectively, diagnoses from 10,825 clinical visits (as of Nov. 2022) of 1,877 participants were considered as the binary phenotype, with 4,010 clinical visits with a cognitively nonimpaired (i.e., normal) diagnosis and 6,815 visits with a cognitively impaired diagnosis (including diagnoses of mild cognitive impairment (MCI) and dementia). The average age at each clinical diagnostic visit was 75.78±6.68 and 75.75±7.82 for nonimpaired (coded as one) and impaired (coded as zero) groups, respectively (Table 1).

#### Longitudinal continuous phenotype: Florbetapir (amyloid) PET-SUVR

A subset of 1,096 participants were subjected to at least one amyloid PET scan with the florbetapir (^18^F) ligand. The composite standardized uptake value ratios (SUVRs), normalized to the cerebellum, for a total of 2,598 longitudinal scans were considered as the continuous phenotype in the following analyses (Table 1). In addition, participants’ amyloid positivity status was determined from the composite SUVR of their latest Florbetapir scan, with a cut-off value of 1.11[23].

### 2.2 Longitudinal GWAS model

#### Varying coefficient model

RVMMAT was applied to genome-wide SNPs in ADNI samples to detect time-varying genetic effects on the longitudinal binary and continuous phenotypes. In RVMMAT, a dynamic genetic effect is modeled using smoothing splines and parameters are estimated by maximizing a double penalized quasi-likelihood function. A joint test is then conducted to combine the *p-values* of two score tests for a fixed genetic effect and a random genetic effect, using a Cauchy combination method. Details about the RVMMAT model formulation and solution can be found in Xu et al., 2022[11].

Besides modeling varying coefficients and using score tests, RVMMAT is a retrospective test in which genotypes are treated as responses conditional on phenotypes and covariates. This strategy enhances robustness against model misspecification and also improves statistical power[19, 24–26]. RVMMAT can be viewed as an extension of the generalized linear mixed model (GLMM) [27] and retrospective GLMM-based association test (RGMMAT) [19] to a varying coefficient mixed effects model [28]. By embracing more flexible assumptions on the genetic effect function, unlike methods that assume constant genetic effects, RVMMAT can detect time-varying genetic variants linked with dynamic phenotypes, resulting in increased statistical power.

#### Applying to ADNI samples

We performed longitudinal GWAS on AD using RVMMAT with cubic smoothing splines in ADNI participants for both binary and continuous phenotypes detailed in section 2.1. Subjects’ sex and the first five principal components of the genetic relationship matrix based on imputed whole-genome genotypes were included as covariates. Age at each diagnosis for all participants were normalized to a range of zero to one and coded as the time factor in RVMMAT.

For comparison, we also ran a longitudinal GWAS model assuming constant genetic effect over time for the same phenotypes on ADNI participants. To this end, RGMMAT[19] was applied with the same set of covariates.

### 2.3 Post hoc analyses on significant SNPs

#### Pathway analyses

We performed pathway enrichment analyses using MetaCore^TM^ on significant SNPs identified by RVMMAT to characterize variants with potential time-varying genetic effects in AD. Significant SNPs were annotated to the genes that they were located within, or to their nearest genes. The obtained SNP and gene lists were input to the functional pathway analyses. Fisher’s exact test was used to test whether SNPs and genes were significantly enriched for a functional pathway.

#### Analysis of time-varying genetic effects

We next examined genetic effects at each time point (i.e., within each age interval) for significant SNPs identified by RVMMAT. All longitudinal visits were grouped into various age ranges with a five-year interval. For each age interval, phenotype and covariates at each longitudinal visit, and corresponding participants’ genotype were obtained. Note that only one visit was kept from the same subject with the same phenotype in each age interval.

We analyzed the time-varying genetic effects on phenotypes using both chi-square (χ^2^) statistics and regression models. For the binary phenotype, within each age interval, we first performed a χ^2^ test to examine the genotypic differences between phenotype groups for each significant SNP. A larger χ^2^ value indicates a greater genetic difference between phenotype groups for the SNP within this age interval, as compared to other age intervals. For both binary and continuous phenotypes, within each age interval, we further performed a logistic regression and a linear regression with genotype as predictor and phenotype as outcome, respectively. The same set of covariates as in RVMMAT were included in the regression analyses. A larger regression coefficient (in amplitude) of genotype for a given age interval indicates a greater genetic effect for that SNP, as compared to other age intervals.

### 2.4 Replication analyses

NACC participants were utilized as a replication dataset. The NACC has been established in collaboration with more than 42 previous and current Alzheimer’s Disease Research Centers (ADRCs) throughout the US over more than 20 years[29].

#### Whole-genome sequencing data

We included data from 785 NACC participants with whole-genome sequencing (WGS) data stored on the National Institute on Aging Genetics of Alzheimer’s Disease Data Storage Site Data Sharing Service (NIAGAD-dss, https://dss.niagads.org/datasets/ng00067/). Data in genome variant calling format (gVCF) were downloaded from the NIAGAD-dss and were aligned to genome reference consortium human build 38 (GRCh38).

#### Binary phenotype

Participants’ sex, race, clinical diagnoses, and age at each clinical diagnosis were obtained from the NACC Uniform Data Set[30], and listed in Table 1. Among the 785 participants, 310 were men and 475 were women; 185 were African Americans (23.57%), 593 were European Americans (75.54%), and 7 were of other races (0.89%). The NACC data has more diverse race distribution as compared to the ADNI participants.

A total of 6,494 clinical visits with a diagnosis of cognitively normal, self-reported impairment (not MCI), MCI, or AD, were obtained for these 785 participants. We again coded each diagnosis into a nonimpaired (cognitively normal, N_visit_=4,708) and an impaired (self-reported impairment, MCI, and AD, N_visit_=1,786) group. The average age at each clinical diagnosis was slightly older in NACC participants, as compared to ADNI participants, with 77.19±7.43 for nonimpaired group and 79.28±10.06 for impaired group (Table 1).

#### RVMMAT

We performed RVMMAT on significant SNPs identified from ADNI data to examine whether the observed time-varying genetic effect on AD could be replicated using the NACC dataset. Since ADNI and NACC data were aligned to different human reference genome versions, we first performed a genome lift-over (https://genome.ucsc.edu/cgi-bin/hgLiftOver) from GRCh37 to GRCh38 for RVMMAT significant SNPs. Corresponding SNPs from NACC participants were then extracted from the gVCF files based on the aligned positions. The same set of covariates were included in the replication analyses.

## 3. Results

### 3.1. Longitudinal GWAS with the binary phenotype

#### 3.1.1 Significant SNPs

In ADNI participants, RVMMAT analysis of the clinical impairment status as a phenotype showed no evidence of inflation in the quantile-quantile plot (Fig. 2(A)), with a genomic inflation factor (-1) of 0.97. RVMMAT significance levels (raw p-values(*p_raw_*)) for all SNPs were shown in a Manhattan plot (Fig. 2(B)). For each chromosome, SNPs with a *p_raw_*<1E-11 were clustered and represented by one triangle at *p_raw_*=1E-11 (solid grey line in Fig. 2(B)). Although we performed genome-wide tests across 9,573,130 imputed SNPs, many SNPs were highly correlated; and therefore, we considered a widely used threshold *p_raw_*<5E-08 as our genome-wide significance level (solid black line in Fig. 2(B)). We further performed a false discovery rate (FDR) correction on *p_raw_*across all SNPs at a lower genome-wide significance level (*p_FDR_*<0.05, dashed black line in Fig. 2(B)).

**Fig. 2.**
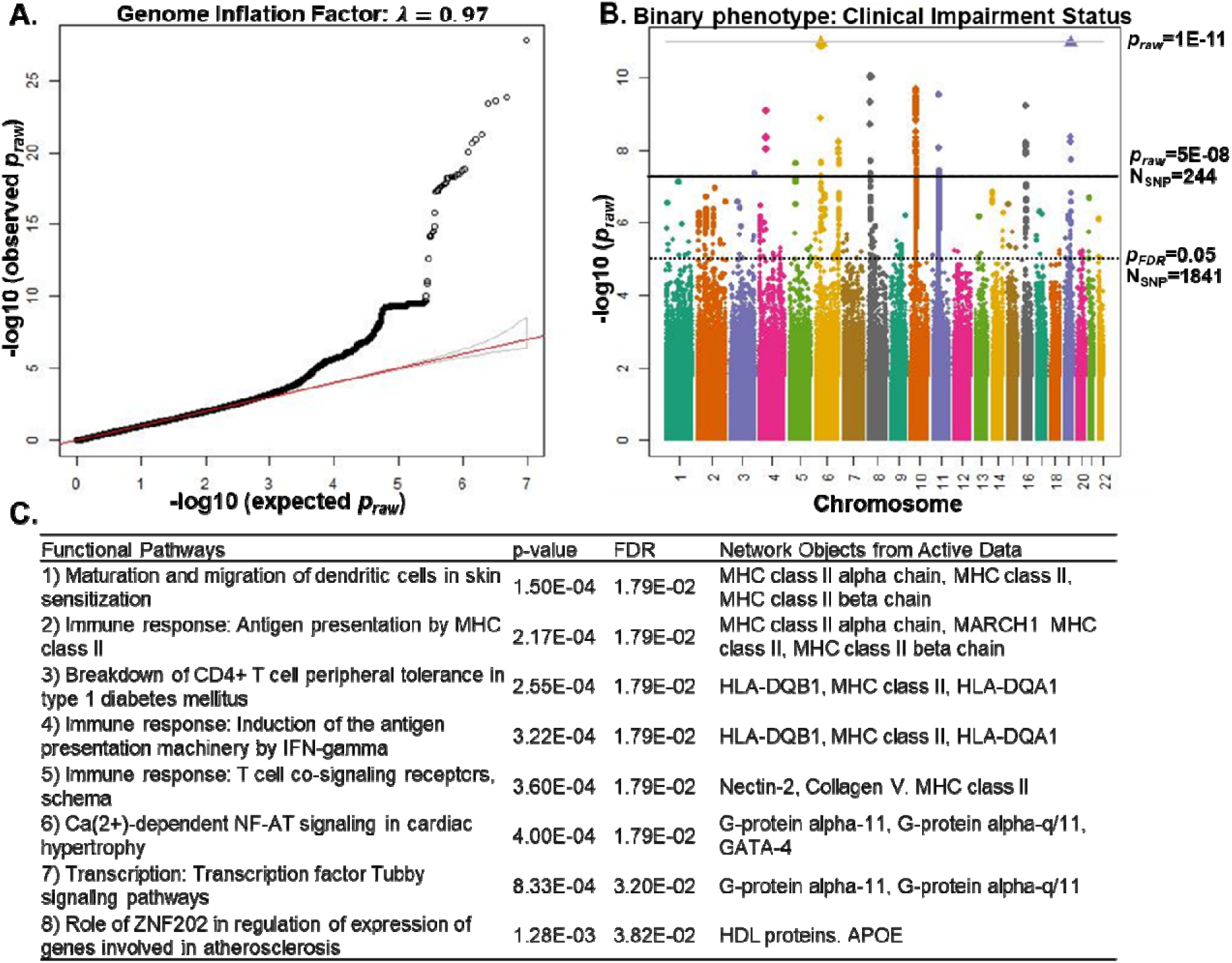
RVMMAT results on binary phenotype of clinical impairment status in AD using ADNI participants. ***(A).*** Quantile-quantile plot scatters the observed *p-values* (y-axis) and the expected *p-values* (x-axis), indicating no evidence of inflation with a genome inflation factor. ***(B).*** Manhattan plot shows that 244 SNPs reached genome-wide significance with *p_raw_*<5E-08 (solid black line), and 1,841 SNPs reached genome-wide significance with *p_FD_* <0.05 (dashed black line). For better display purposes, SNPs with a *p_raw_*<1E-11 (solid grey line) were not shown individually and were clustered and represented by one triangle at *p_raw_*=1E-11 (solid grey line). ***(C).*** Significantly disrupted functional pathways on 1,841 genome-wide significant SNPs with *p_FDR_* <0.05 (and their annotated genes) using MetaCore^TM^.

At *p_raw_*<5E-08, 244 SNPs reached genome-wide significance (Fig. 2(B)). The SNP genome positions, RVMMAT statistics (*p_raw_* and *p_FDR_*), minor allele frequencies (MAFs), annotated genes, and distances to annotated genes (in base-pairs) are listed in Supplement Table 1. Among the 244 SNPs, the most significant signals were derived from 36 SNPs on chromosome 19, and were clustered at the *APOE, TOMM40, APOC1* and *NECTIN2* (also known as *PVRL2*) genes. Among which, the most significant SNP was rs429358 at position 45411941 on chromosome 19 (*p_raw_*=1.66E-28), which was the *APOE* E4 determinant SNP. Fourteen SNPs on chromosome 6 were clustered at the *HLA-DQB1, HLA-DQA1,* and *TCP10* genes (smallest *p_raw_*=8.48E-12); 35 SNPs on chromosome 10 were located at the *LINC00999* gene (smallest *p_raw_*=2.23E-10); six SNPs on chromosome 4 were located at the *SLAIN2* gene (smallest *p_raw_*=8.23E-10), and one SNP on chromosome 5 was located at the *EMB* gene (*p_raw_*=2.28E-08). Besides these protein-coding genes, 131 SNPs on chromosome 10, eight SNPs on chromosome 16, six SNPs on chromosome 8, and six SNPs on chromosome 11 were located at *HSD17B7P2, SEPT7P9, KSR1P1, HERC2P4, RPS3AP35* and *OR4A49P* pseudogenes or other non-protein coding genes (Supplement Table 1).

We next examined functional pathways associated with the 1,841 genome-wide significant SNPs at *p_FDR_*<0.05 (Fig. 2(C) and Supplement Table 2). Considering a Fisher’s exact test for the enrichment statistics, we identified eight significantly enriched functional pathways (FDR-p<0.05 in Fisher’s exact test) associated with immune response (four pathways), G-protein signaling (two pathways), lipid-associated gene expression (one pathway) and dendritic cell maturation (one pathway).

#### 3.1.2. Time-varying genetic effect

The genotypic differences and the estimated genetic effect over time are shown in Fig. 3 for 244 genome-wide significant SNPs at *p_raw_*<5E-08. The χ^2^ statistics (Fig. 3(A)) and estimated logistic regression coefficients (Fig. 3(B)) at each time point (i.e., age interval) were obtained by using the observed phenotypic values within that age range. A straight line was used to connect the estimated values at two adjacent age intervals, and a zero-line was added to indicate no genotypic effect in Fig. 3(B).

**Fig. 3:**
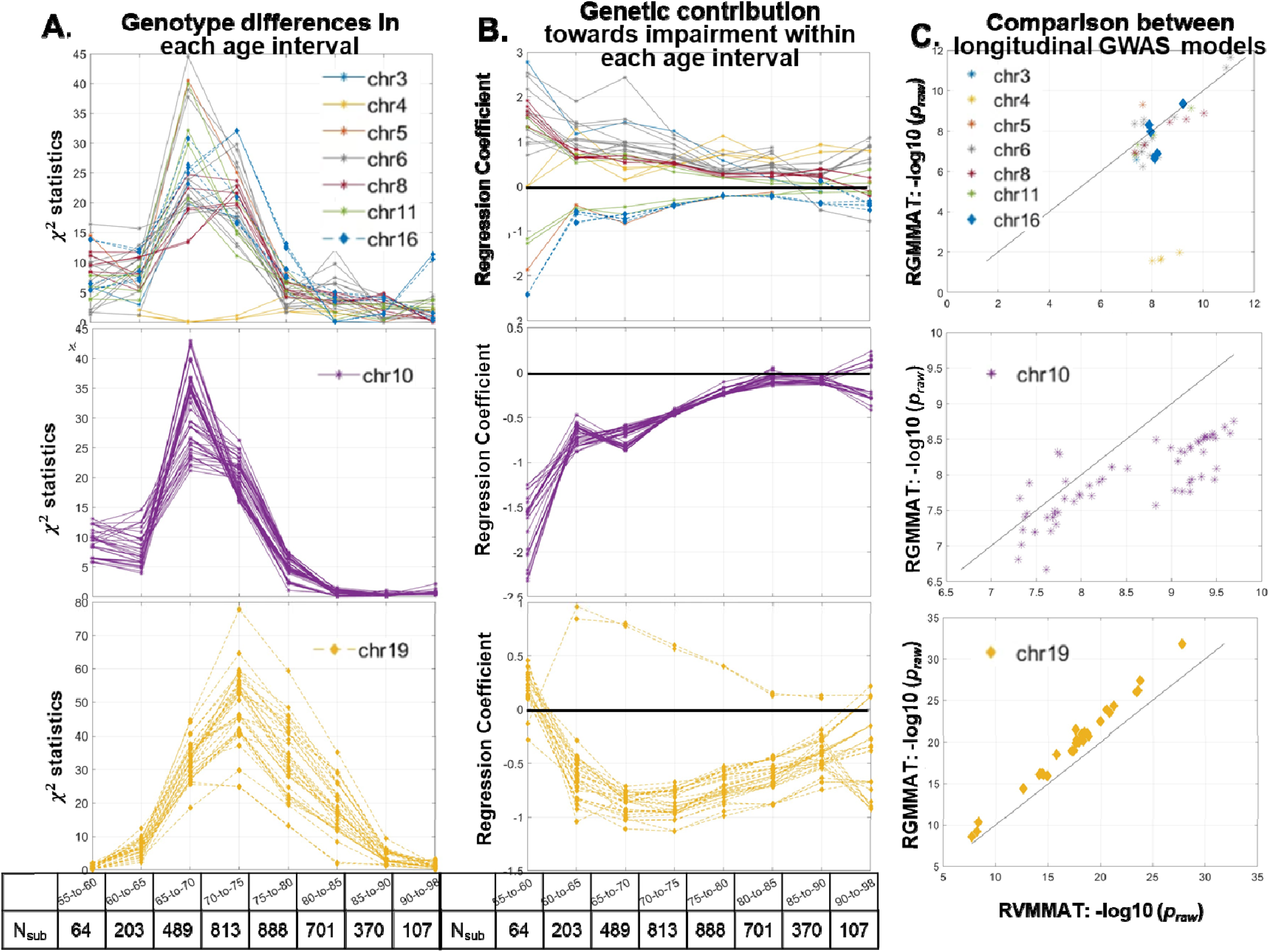
Estimated genotypic difference and genetic effect of 244 genome-wide significant SNPs at *p_RAW_* <5E-08 on binary clinical impairment status in AD at each time point and age interval. SNPs on chromosomes 10 and 19 are plotted separately in the middle (chromosome 10 SNPs) and bottom panels (chromosome 19 SNPs) due to the relatively larger number of significant SNPs. SNPs on other chromosomes are plotted in the top panel. *(A).* Estimated genotypic differences (χ^2^ statistics) between phenotype groups within each age interval. Number of subjects in each age group are listed at the bottom. *(B).* Estimated genotypic effect in predicting impairment status (regression coefficient) in logistic regression model for subjects within each age interval. Deviations from the zero-line (i.e., no genotypic effect (solid black line)) indicate greater genotypic effects on this phenotype. *(C)*. Comparisons of significance levels (-log10(*p_raw_*)) between longitudinal GWAS models with time-varying coefficients (RVMMAT, x-axis) and assuming time-constant genetic effect (RGMMAT, y-axis).

As shown in Fig. 3(A), for SNPs on chromosome 19, we observed an overall greater genotypic difference between clinically impaired and nonimpaired participants in a wider age interval of 65-80 years old (maximum χ^2^=77.86, bottom plot in Fig. 3(A)), as compared to other SNPs (top and middle plots in Fig. 3(A)). An overall greater genotypic effect on this phenotype was also observed in the same age interval for these SNPs, as reflected by larger (in amplitude) logistic regression coefficients that deviate from the zero-line (Fig. 3(B), bottom plot). Since in the logistic regression model, we coded clinically impaired status as zero and clinically nonimpaired status as one, a negative regression coefficient here indicated the possession of alleles contributing negatively towards clinically normal status, (i.e., increased effect towards clinical impairment status). Furthermore, for these chromosome 19 SNPs, both longitudinal GWAS models with and without time varying coefficient (RVMMAT and RGMMAT) demonstrated high statistical power, as most SNPs reached significant *p-values* (*p_raw_*<1E-10) in both models (Fig. 3(C), bottom plots).

We additionally observed a larger genotypic difference on AD clinical impairment status in a smaller age interval of 70-75 years old for SNPs on chromosomes 6, 8, 10 and 11 (top and middle plots in Fig. 3(A)). These genotypic differences were not observed before 60 years old, and diminished after 80 years old, as shown by the reduced x^2^ values. Deviations of regression coefficients from the zero-line in Fig. 3(B) also indicated that the genetic effect of these SNPs on AD clinical impairment status decreased during aging (top and middle plots in Fig. 3(B)). For most of these SNPs, longitudinal GWAS models assuming time-constant genetic effect (RGMMAT) lost power and generated less significant *p-values*, as compared to the model with time-varying coefficient (RVMMAT, Fig. 3(C) upper and middle plots).

#### 3.1.3 Replication

We repeated the RVMMAT method on 244 genome-wide significant SNPs (*p_raw_*<5E-08) using the NACC dataset. In this replication, 29 SNPs reached *p_raw_*<0.05 (Supplement Table 1). Among which, 24 SNPs were clustered *at APOE, TOMM40, NECTIN2*, and *APOC1* genes on chromosome 19, and five SNPs were clustered at *KSR1P1* pseudogene on chromosome 10. Two additional SNPs clustered to *HLA-DQB1* gene on chromosome 6 reached a trend level significance (*p_raw_* <0.10).

We further performed an FDR correction on these raw *p-values* to correct for multiple comparisons. In this correction, we focused on SNPs with similar MAFs between the ADNI and NACC participants (N_SNP_ = 194, detailed in Supplement Material 1), and 12 SNPs on chromosome 19 survived this FDR-correction (Supplement Fig. 1).

### 3.2 Longitudinal GWAS with a continuous phenotype

Fig. 4 shows RVMAMT results on a continuous phenotype of brain amyloid accumulation using 2,598 longitudinal PET-SUVR measures from ADNI participant. Using this phenotype, RVMMAT produced a genomic inflation factor of 0.95 (Fig. 4(A)). At *p_raw_*<5E-08, 73 SNPs reached genome-wide significance (Fig. 4(B)). Most of these SNPs were again clustered on chromosome 19 and annotated to *APOE, APOC1, TOMM40* and *NECTIN2* genes. Besides chromosome 19, one SNP on chromosome 1 reached genome-wide significance and was located at the *FMN2* gene. In addition, as shown in Fig. 4(C), an increasing genotypic effect on PET-SUVR was observed when participants’ age increased, as indicated by the larger (in amplitude) regression coefficient that deviates from the zero-line. (don’t have a para separation here). We next performed a functional pathway analysis using top SNPs with *p_raw_*<1E-04 (N_SNP_=1,039, Supplement Table 4) and identified 17 significantly disrupted biological processes that were involved in immune response, signal transduction, development, and neurophysiological process (Supplement Table 5).

**Fig. 4.**
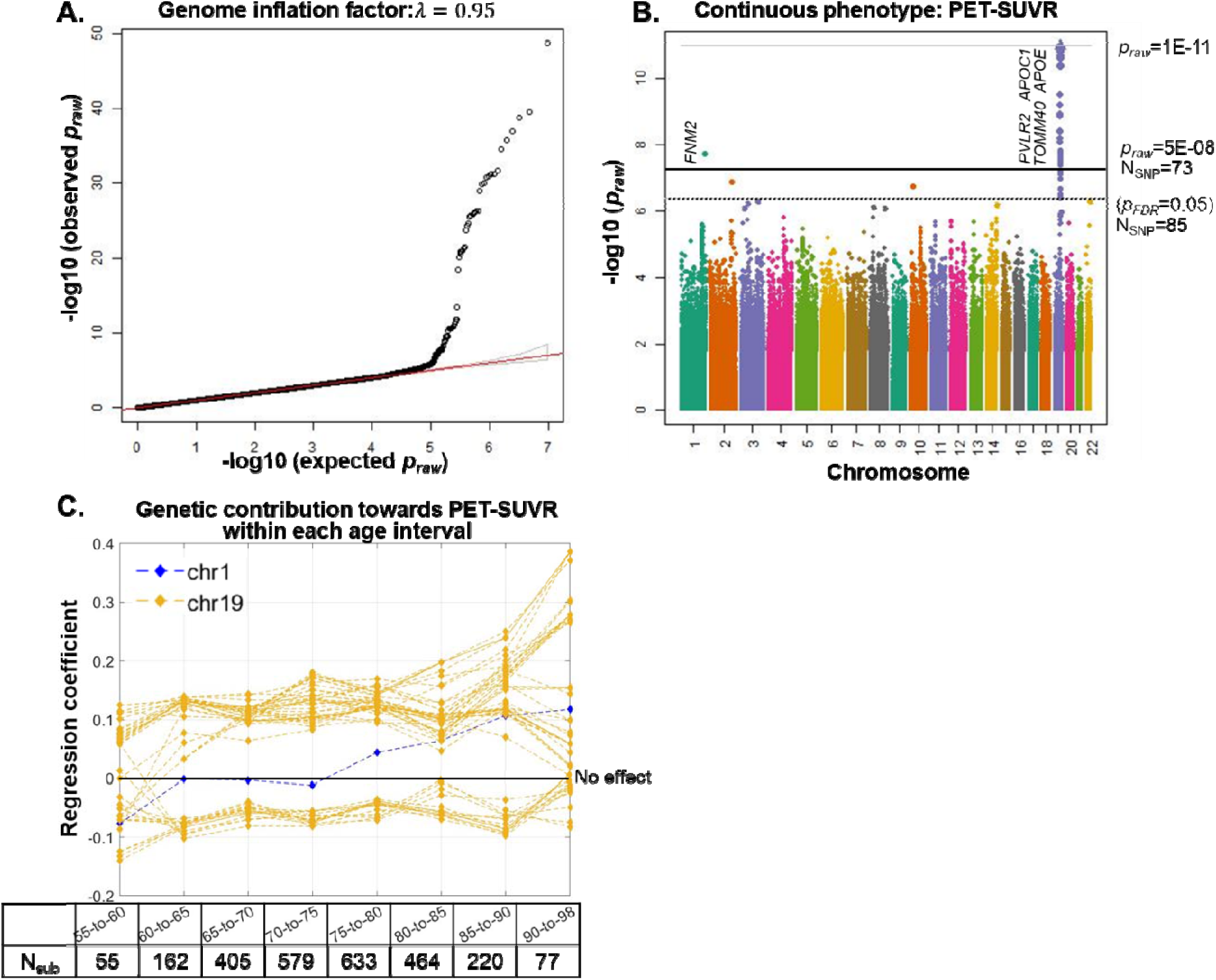
RVMMAT results on continuous phenotype of brain amyloid accumulation (PET-SUVR) in AD using ADNI participants. *(A).* Quantile-quantile plot indicates no evidence of inflation with a genome inflation factor =0.95. *(B).* Manhattan plot shows that 73 SNPs reached genome-wide significance with *p_raw_*<5E-08 (solid black line), and 85 SNPs reached genome-wide significance with *p_FDR_*<0.05 (dashed black line). For better display purposes, SNPs with a *p_raw_*<1E-11 (solid grey line) were not shown individually and were clustered and represented by one triangle at *p_raw_*=1E-11 (solid grey line). *(C).* Estimated genotypic effect (regression coefficient) in predicting PET-SUVRs in linear regression model for subjects within each age interval. Deviations from the zero-line (solid black line) indicate greater genotypic effects on this phenotype.

## 4. Discussion

The development and progression of AD is a complex process that could change over time, during which the impact of genetic variation on phenotypes may also fluctuate. Incorporating longitudinal phenotypes with time-varying coefficients in GWAS provides the opportunity to identify this changing genetic effect on phenotypic variations over time along the disease continuum; and therefore, may shed new light on understanding AD patho-mechanisms. In this study, we utilized a varying coefficient model, RVMMAT, to perform longitudinal GWAS on repeated measurements of clinical and biological phenotypes in AD. Benefiting from the improved statistical power, a relatively limited number of samples from the publicly available ADNI database were used. Our study led to the identification of genome-wide significant SNPs that could convey time-varying and age-dependent genetic effects on phenotypes for both binary clinical impairment status and continuous amyloid accumulation in AD.

Modeling varying AD genetic risk over time has been shown to result in a 5-10% statistical power gain in GWAS and has assisted in discovering novel AD-associated variants[17, 20, 31]. The method we adopted here, RVMMAT, utilized time-varying coefficients to model a fluctuating genetic effect on dynamic phenotypes, and therefore could also lead to the improved statistical power in GWAS. In applying to other complex longitudinal phenotypes, this retrospective time-varying model has been shown to 1) improve the statistical power, 2) be robust to model misspecification, and 3) identify novel genetic variants, as compared to linear mixed models with and without time-varying coefficients[11]. In our study, the RVMMAT method produced significant results for SNPs conveying moderate genetic risks, as compared to models assuming constant genetic effect over time (i.e., RGMMAT, Fig. 3(C), upper and middle plots). Note that longitudinal GWAS methods, both with and without modeling time-varying genetic effect, tend to produce significant results for SNPs that convey relatively large genetic effects (Fig. 3(C) bottom plots).

With increased statistical power using RVMMAT, we identified 244 genome-wide significant SNPs relevant to clinical impairment status in AD (Fig. 2). The most predominant signals were clustered on chromosome 19, with the most significant SNP to be the *APOE* E4 determinant SNP (Supplement table 1). Our post-hoc age-stratified analyses further demonstrated that the maximum genotypic effect of these SNPs on AD impairment status existed at the age interval of 70-75 years old, and then declined with age (Fig. 3). These observations were consistent with several previous reports. For instance, *APOE* E4 allele has been shown to convey an age-dependent hazard in developing AD that declines with increasing age[12]. Logistic regression analyses (similar to what we performed in this study) have further demonstrated that the *APOE* genotype shared a greater regression coefficient and a larger area under the ROC curve in predicting clinical AD cases in participants younger than 80 years old, as compared to more senior participants[32]. In our study, these observed time-varying genetic effects on chromosome 19 SNPs, especially the *APOE* E4 determinant SNP, were also replicated using the NACC dataset (Supplement Table 1). Our replication study combined with previous reports have strengthened the potential time-varying effect of these chromosome 19 SNPs on AD clinical status.

Different from clinical impairment status, our study identified 72 SNPs on chromosome 19 that may convey an increasing genotypic effect on brain amyloid accumulation with increased age (Fig. 4). More specifically, our results showed that before the age of 85, the genotypic effect of these SNPs on brain amyloid accumulation remained constant, with a significant increase in more senior participants. These results indicated that distinct time-varying genetic effects might occur for these chromosome 19 SNPs towards AD clinical and biological phenotypes, respectively. These results will benefit from further validation, as our study contained a relatively small number of senior participants for the amyloid phenotype.

Besides chromosome 19, our longitudinal GWAS models identified significant SNPs on other chromosomes. Among them, SNPs on chromosome 6 were clustered to *HLA-DQB1* gene and reached a trend level significance in our replication analysis using the NACC cohort (Fig. 2 and Supplement Table 1). *HLA-DQB1* has been reported to convey a significant age-of-onset risk in developing AD[33], which might explain our observation of the decreased genotypic effect of these SNPs on AD clinical impairment status along aging. Our study further identified significant SNPs clustered to lincRNA or pseudogenes on chromosome 10, with a time-varying genetic effect on AD clinical impairment status. More recently, there has been an increasing number of studies focused on the role of long non-coding RNAs in regulating expression and modulating protein levels in AD[34, 35]. Therefore, these SNPs could be potential novel signals in AD genetic mechanisms, but require further replication and validation studies.

Our pathway analyses of top SNPs highlighted the involvement of immune responses, lipid metabolism, G-protein signaling and neurophysiological processes in clinical impairment status and brain amyloid accumulations related to AD (Fig. 2 and Supplement Table 3). Given the established role for these pathways in neurodegeneration[7], our findings suggest that the longitudinal GWAS model can provide enhanced statistical power in detecting biologically relevant genetic loci that are associated with phenotypic dynamics, and highlight the role of neuroinflammation in AD.

Following the ATN framework[1] and AD clinical impairment status, we have further limited our longitudinal GWAS analyses to participants that were classified as amyloid positive on their latest PET scans in ADNI (detailed in Supplement Material 2). In this analysis, chromosome 19 SNPs clustered to *APOE, APOC1* or *TOMM40* did not reach genome-wide significance (Supplement Fig. 2). Given the reduced sample size (N_sub_=620), we may lack the statistical power of the model to detect relevant signals. The lack of significance of these chromosome 19 SNPs might also provide additional evidence of previous findings that SNPs clustered to *APOE* loci were associated with amyloid accumulation in AD[36]. Furthermore, pathway analyses on these top SNPs have highlighted oxidative stress, particularly reactive oxygen species-induced cellular signaling, to be the most disrupted functional pathway (Supplement Fig. 2(C)).

## 5. Limitations

Interestingly, we were only able to replicate time-varying genetic signals on chromosome 19 using the NACC dataset for the phenotype of AD clinical impairment status. This limited replication might be explained partially by the different racial and phenotypic group distributions between ADNI and NACC participants (Table 1). Future analyses using additional cohorts from diverse backgrounds could then take full advantage of the RVMMAT method in longitudinal GWAS to produce novel and stable results.

Our study did not lead to the identification of genome-wide significant SNPs associated with several previously identified AD-related genetic risk loci or genes, such as *CR1, CLU, TREM2, ABCA1,* or *SORL1* [4, 7]. The relatively small sample size, even after incorporating longitudinal measures, could contribute to this observation. The potential limited time-varying effect for SNPs associated with these genes could impose additional difficulties to reach genome-wide significance in RVMMAT. Future analyses with a larger sample size and more longitudinal follow-up studies could increase the sensitivity of the RVMMAT method for these previously reported AD-associated genes.

With the larger sample-size, it would also be interesting to restrict longitudinal GWAS, or even traditional GWAS, to biologically defined AD cases, i.e., to amyloid-positive and tau-positive participants. Our current study did not have sufficient participants or statistical power to be restricted to these biological AD cases. This potential future analysis could assist in further elucidating the genetic mechanisms and time-varying genetic contributions in biological AD.

One possible future direction that might not rely on additional participants would be to compute an age-specific polygenic risk score (PRS) for each age-group with the genetic risk estimated particularly for that age interval. PRS has been shown to have a greater impact in developing clinical AD as participants’ age increase[32]. It is interesting that in our analysis on the AD clinical phenotype, only a few genome-wide significant SNPs conveyed an increasing genotypic effect along aging. Age-specific PRS might be a less biased and more accurate measure to estimate participants’ genetic risk, as it accounts for the potential time-varying genotypic effects on dynamic phenotypes.

## 6. Conclusions

In summary, we have tested our hypothesis that the RVMMAT method could be used to detect a time-varying genetic effect on clinical and biological longitudinal phenotypes in AD. Additionally, we have 1) corroborated previous findings of declined phenotypic effects during aging for most SNPs; 2) highlighted the involvement of neuroinflammation in AD; and 3) provided potential novel genetic signals related to AD. Future analyses with a larger and more diverse sample are necessary to replicate and validate our findings.

## 7. List of Abbreviations

AD: Alzheimer’s disease
MCI: Mild cognitive impairment
ADNI: Alzheimer’s Disease Neuroimaging Initiative
NACC: National Alzheimer’s Coordinating Center
ADRC: Alzheimer’s Disease Research Centers
GRCh37/38: Genome Reference Consortium Human Build 37/38
SNP: Single Nucleotide Polymorphism
GWAS: Genome Wide Association Studies
WGS: Whole-genome Sequencing
PET: Positron Emission Tomography
SUVR: Standardized Uptake Value Ratio
NIAGADS: National Institute on Aging Genetics of Alzheimer’s Disease Data Storage site
RVMMAT: Retrospective Varying coefficient Mixed Model Association Test
RGMMAT: Retrospective Generalized linear Mixed Model Association Test
FWE: Bonferroni based Family Wise Error rate correction for multiple comparison
FDR: False Discovery Rate based correction for multiple comparison
gVCF: genome Variant Calling Files

## 8. Declaration

### Ethics approval and consent to participate

#### ADNI participants

Main data used in this study are collected by ADNI. Therefore, the study was approved by each participating ADNI site’s local Institutional Review Boards, as documented on the ADNI website. All participants gave written, informed consent. No ethics approval or participant consent was obtained locally for ADNI participants.

#### NACC participants

Replication data used in this study are collected by each ADRC and uploaded to the NACC database. Therefore, the study was approved by each ADRC site’s local Institutional Review Boards. No ethics approval or participant consent was obtained locally for NACC participants.

## Consent for publication

Not applicable.

## Availability of data and materials

ADNI participants used in this study are 1,877 individuals with whole-genome genotyping data available on https://ida.loni.usc.edu/pages/access/geneticData.jsp through ADNI data request process. De-identified data from NACC participants used in this study are available through NACC data request process (https://naccdata.org/requesting-data/data-request-process) with qualified researchers.

## Competing interests

All authors declare that the research was conducted in the absence of any commercial or financial relationships that could be construed as a potential conflict of interest.

## Authors’ contributions

**Xiaowei Zhuang:** Conceptualization, Methodology, Formal analysis, Investigation, Visualization, Writing original draft, Review & editing. **Gang Xu:** Conceptualization, Methodology, Formal analysis, Investigation, Visualization, Writing original draft, Review & editing. **Zuoheng Wang:** Methodology, Visualization, Review & editing. **Dietmar Cordes:** Visualization, Review & editing. **Amei Amei:** Visualization, Review & editing. **Edwin C. Oh:** Conceptualization, Methodology, Formal analysis, Investigation, Visualization, Infrastructure provision, Review & editing, Supervision.

## Funding

Dr. Oh is supported by an NIH grant R01-MH109706. Dr. Xu and Dr. Wang are supported by the NIH grant R01LM014087. Ms. Zhuang and Dr. Cordes are supported by the NIH grant RF1-AG071566.

## Acknowledgements

Data collection and sharing for this project was funded by the Alzheimer’s Disease Neuroimaging Initiative (ADNI) (National Institutes of Health Grant U01 AG024904) and DOD ADNI (Department of Defense award number W81XWH-12-2-0012). ADNI is funded by the National Institute on Aging, the National Institute of Biomedical Imaging and Bioengineering, and through generous contributions from the following: AbbVie, Alzheimer’s Association; Alzheimer’s Drug Discovery Foundation; Araclon Biotech; BioClinica, Inc.; Biogen; Bristol-Myers Squibb Company; CereSpir, Inc.; Cogstate; Eisai Inc.; Elan Pharmaceuticals, Inc.; Eli Lilly and Company; EuroImmun; F. Hoffmann-La Roche Ltd and its affiliated company Genentech, Inc.; Fujirebio; GE Healthcare; IXICO Ltd.; Janssen Alzheimer Immunotherapy Research & Development, LLC.; Johnson & Johnson Pharmaceutical Research & Development LLC.; Lumosity; Lundbeck; Merck & Co., Inc.; Meso Scale Diagnostics, LLC.; NeuroRx Research; Neurotrack Technologies; Novartis Pharmaceuticals Corporation; Pfizer Inc.; Piramal Imaging; Servier; Takeda Pharmaceutical Company; and Transition Therapeutics. The Canadian Institutes of Health Research is providing funds to support ADNI clinical sites in Canada. Private sector contributions are facilitated by the Foundation for the National Institutes of Health (www.fnih.org). The grantee organization is the Northern California Institute for Research and Education, and the study is coordinated by the Alzheimer’s Therapeutic Research Institute at the University of Southern California. ADNI data are disseminated by the Laboratory for Neuro Imaging at the University of Southern California.

The NACC database is funded by NIA/NIH Grant U24 AG072122. NACC data are contributed by the NIA-funded ADRCs: P30 AG062429 (PI James Brewer, MD, PhD), P30 AG066468 (PI Oscar Lopez, MD), P30 AG062421 (PI Bradley Hyman, MD, PhD), P30 AG066509 (PI Thomas Grabowski, MD), P30 AG066514 (PI Mary Sano, PhD), P30 AG066530 (PI Helena Chui, MD), P30 AG066507 (PI Marilyn Albert, PhD), P30 AG066444 (PI John Morris, MD), P30 AG066518 (PI Jeffrey Kaye, MD), P30 AG066512 (PI Thomas Wisniewski, MD), P30 AG066462 (PI Scott Small, MD), P30 AG072979 (PI David Wolk, MD), P30 AG072972 (PI Charles DeCarli, MD), P30 AG072976 (PI Andrew Saykin, PsyD), P30 AG072975 (PI David Bennett, MD), P30 AG072978 (PI Neil Kowall, MD), P30 AG072977 (PI Robert Vassar, PhD), P30 AG066519 (PI Frank LaFerla, PhD), P30 AG062677 (PI Ronald Petersen, MD, PhD), P30 AG079280 (PI Eric Reiman, MD), P30 AG062422 (PI Gil Rabinovici, MD), P30 AG066511 (PI Allan Levey, MD, PhD), P30 AG072946 (PI Linda Van Eldik, PhD), P30 AG062715 (PI Sanjay Asthana, MD, FRCP), P30 AG072973 (PI Russell Swerdlow, MD), P30 AG066506 (PI Todd Golde, MD, PhD), P30 AG066508 (PI Stephen Strittmatter, MD, PhD), P30 AG066515 (PI Victor Henderson, MD, MS), P30 AG072947 (PI Suzanne Craft, PhD), P30 AG072931 (PI Henry Paulson, MD, PhD), P30 AG066546 (PI Sudha Seshadri, MD), P20 AG068024 (PI Erik Roberson, MD, PhD), P20 AG068053 (PI Justin Miller, PhD), P20 AG068077 (PI Gary Rosenberg, MD), P20 AG068082 (PI Angela Jefferson, PhD), P30 AG072958 (PI Heather Whitson, MD), P30 AG072959 (PI James Leverenz, MD).

The authors additionally thank all NACC and ADNI participants and their families for their support in AD research.

## Notes

### Competing Interest Statement

The authors have declared no competing interest.

## References

1. Jack CR, Bennett DA, Blennow K, Carrillo MC, Dunn B, Haeberlein SB, et al. NIA-AA Research Framework: Toward a biological definition of Alzheimer’s disease. Alzheimer’s and Dementia. 2018;14:535–62.

2. Heneka MT, Carson MJ, Khoury J El, Landreth GE, Brosseron F, Feinstein DL, et al. Neuroinflammation in Alzheimer’s disease. Lancet Neurol. 2015;14:388–405.

3. Gatz M, Reynolds CA, Fratiglioni L, Johansson B, Mortimer JA, Berg S, et al. Role of genes and environments for explaining Alzheimer disease. Arch Gen Psychiatry. 2006;63:168–74.

4. Lambert JC, Ibrahim-Verbaas CA, Harold D, Naj AC, Sims R, Bellenguez C, et al. Meta-analysis of 74,046 individuals identifies 11 new susceptibility loci for Alzheimer’s disease. Nat Genet. 2013;45:1452–8.

5. Wightman DP, Jansen IE, Savage JE, Shadrin AA, Bahrami S, Holland D, et al. A genome-wide association study with 1,126,563 individuals identifies new risk loci for Alzheimer’s disease. Nat Genet. 2021;53:1276–82.

6. Kunkle BW, Grenier-Boley B, Sims R, Bis JC, Damotte V, Naj AC, et al. Genetic meta-analysis of diagnosed Alzheimer’s disease identifies new risk loci and implicates Aβ, tau, immunity and lipid processing. Nat Genet. 2019;51:414–30.

7. Bellenguez C, Küçükali F, Jansen IE, Kleineidam L, Moreno-Grau S, Amin N, et al. New insights into the genetic etiology of Alzheimer’s disease and related dementias. Nat Genet. 2022;54:412–36.

8. Jansen IE, Savage JE, Watanabe K, Bryois J, Williams DM, Steinberg S, et al. Genome-wide meta-analysis identifies new loci and functional pathways influencing Alzheimer’s disease risk. Nat Genet. 2019;51:404–13.

9. Bryois J, Buil A, Ferreira PG, Panousis NI, Brown AA, Viñuela A, et al. Time-dependent genetic effects on gene expression implicate aging processes. Genome Res. 2017;27:545–52.

10. Chu W, Li R, Liu J, Reimherr M. Feature selection for generalized varying coefficient mixed-effect models with application to obesity gwas. Annals of Applied Statistics. 2020;14:276–98.

11. Xu G, Amei A, Wu W, Liu Y, Shen L, Oh EC, et al. Retrospective varying coefficient association analysis of longitudinal binary traits: application to the identification of genetic loci associated with hypertension. bioRxiv. 2022;:2022.10.31.514543.

12. Liu L, Caselli RJ. Age stratification corrects bias in estimated hazard of APOE genotype for Alzheimer’s disease. Alzheimer’s and Dementia: Translational Research and Clinical Interventions. 2018;4:602–8.

13. Petersen RC, Aisen PS, Beckett LA, Donohue MC, Gamst AC, Harvey DJ, et al. Alzheimer’s Disease Neuroimaging Initiative (ADNI): Clinical characterization. Neurology. 2010;74:201.

14. Beekly DL, Ramos EM, van Belle G, Deitrich W, Clark AD, Jacka ME, et al. The National Alzheimer’s Coordinating Center (NACC) Database An Alzheimer Disease Database Purpose of the Database. Alzheimer Dis Assoc Disord. 2004;18:270.

15. Sudlow C, Gallacher J, Allen N, Beral V, Burton P, Danesh J, et al. UK Biobank: An Open Access Resource for Identifying the Causes of a Wide Range of Complex Diseases of Middle and Old Age. PLoS Med. 2015;12:1001779.

16. Bolker BM. Linear and generalized linear mixed models. 2015.

17. Phillips J, Dumitrescu L, Archer DB, Smith AN, Mukherjee S, Lee ML, et al. Longitudinal GWAS Identifies Novel Genetic Variants and Complex Traits Associated with Resilience to Alzheimer’s Disease. Alzheimer’s & Dementia. 2022;18.

18. Yuan M, Xu XS, Yang Y, Zhou Y, Li Y, Xu J, et al. SCEBE: An efficient and scalable algorithm for genome-wide association studies on longitudinal outcomes with mixed-effects modeling. Brief Bioinform. 2021;22.

19. Wu W, Wang Z, Xu K, Zhang X, Amei A, Gelernter J, et al. Retrospective association analysis of longitudinal binary traits identifies important loci and pathways in cocaine use. Genetics. 2019;213:1225–36.

20. Li Y, Xu I, Liu C. Functional Data Modeling and Hypothesis Testing for Longitudinal Alzheimer Genome-Wide Association Studies. 2021. p. 353–79.

21. Saykin AJ, Shen L, Yao X, Kim S, Nho K, Risacher SL, et al. Genetic studies of quantitative MCI and AD phenotypes in ADNI: Progress, opportunities, and plans. Alzheimer’s and Dementia. 2015;11:792–814.

22. Das S, Forer L, Schönherr S, Sidore C, Locke AE, Kwong A, et al. Next-generation genotype imputation service and methods. Nat Genet. 2016;48:1284–7.

23. Landau SM, Fero A, Baker SL, Koeppe R, Mintun M, Chen K, et al. Measurement of Longitudinal β-Amyloid Change with 18F-Florbetapir PET and Standardized Uptake Value Ratios. Journal of Nuclear Medicine. 2015;56:567–74.

24. Hayeck TJ, Zaitlen NA, Loh PR, Vilhjalmsson B, Pollack S, Gusev A, et al. Mixed model with correction for case-control ascertainment increases association power. Am J Hum Genet. 2015;96:720–30.

25. Jiang D, Mbatchou J, McPeek MS. Retrospective Association Analysis of Binary Traits: Overcoming Some Limitations of the Additive Polygenic Model. Hum Hered. 2015;80:187.

26. Wu X, McPeek MS. L-GATOR: Genetic Association Testing for a Longitudinally Measured Quantitative Trait in Samples with Related Individuals. Am J Hum Genet. 2018;102:574.

27. Chen H, Wang C, Conomos MP, Stilp AM, Li Z, Sofer T, et al. Control for Population Structure and Relatedness for Binary Traits in Genetic Association Studies via Logistic Mixed Models. Am J Hum Genet. 2016;98:653–66.

28. Zhang D. Generalized Linear Mixed Models with Varying Coefficients for Longitudinal Data. Biometrics. 2004;60:8–15.

29. Beekly D, Schwabe-Fry K, Bollenbeck M, Thomas G, DeCarli CS, Carmichael OT, et al. [P1–405]: THE NATIONAL ALZHEIMER’S COORDINATING CENTER: DEVELOPMENT OF THE MRI, PET AND CSF BIOMARKER DATABASES. Alzheimer’s & Dementia. 2017;13 7S_Part_8.

30. Besser L, Kukull W, Knopman DS, Chui H, Galasko D, Weintraub S, et al. Version 3 of the National Alzheimer’s Coordinating Center’s Uniform Data Set. Alzheimer Dis Assoc Disord. 2018.

31. Le Guen Y, Belloy ME, Napolioni V, Eger SJ, Kennedy G, Tao R, et al. A novel age-informed approach for genetic association analysis in Alzheimer’s disease. Alzheimers Res Ther. 2021;13.

32. Bellou E, Baker E, Leonenko G, Bracher-Smith M, Daunt P, Menzies G, et al. Age-dependent effect of APOE and polygenic component on Alzheimer’s disease. Neurobiol Aging. 2020;93:69–77.

33. Desikan RS, Fan CC, Wang Y, Schork AJ, Cabral HJ, Cupples LA, et al. Genetic assessment of age-associated Alzheimer disease risk: Development and validation of a polygenic hazard score. PLoS Med. 2017;14:1–17.

34. Shukla S, Srividya K, Nazir A. Not a piece of junk anymore: Pseudogene T04B2.1 performs non-conventional regulatory role and modulates aggregation of α-synuclein and β-amyloid proteins in C. elegans. Biochem Biophys Res Commun. 2021;539:8–14.

35. Li D, Zhang J, Li X, Chen Y, Yu F, Liu Q. Insights into lncRNAs in Alzheimer’s disease mechanisms. RNA Biol. 2021;18:1–11.

36. Morris JC, Roe CM, Xiong C, Fagan AM, Goate AM, Holtzman DM, et al. APOE predicts amyloid-beta but not tau Alzheimer pathology in cognitively normal aging. Ann Neurol. 2010;67:122–31.

