## Supplement Material for "Detecting time-varying genetic effects in Alzheimer’s disease using a longitudinal GWAS model"

**Supplement Materials**

### Supplement Material 1. Replication analyses

We repeated the RVMMAT method on 244 genome-wide significant SNPs (*p_raw_*<5E-08 in ADNI participants) using NACC participants. Supplement Fig. 1(A) below plots the minor allele frequencies (MAFs) of these 244 SNPs in both ADNI (x-axis) and NACC (y-axis) databases. As we stated in the main text, in this replication, 29 out of 244 SNPs demonstrated a *p_raw_*<0.05 (Supplement Table 1). We further performed an FDR correction on these raw p-values to correct for multiple comparisons.

As shown in Supplement Fig. 1(A), for some SNPs, the MAFs differed significantly between the ADNI and NACC participants. We suspected that it might be due to the different racial distributions between two cohorts, and the different sequencing methods in ADNI (genotyping + imputation) and NACC (whole-genome-sequencing). Therefore, in the FDR correction, we focused on 194 SNPs with a consistent MAFs in both databases (i.e., MAF differences less than 10% between these 2 databases; Supplement Fig. 1(B)). In this case, 12 out of these 29 SNPs on chromosome 19 with *p_raw_*<0.05 retained to be significant at *p_FDR_*<0.05 (Fig. 1(C)).

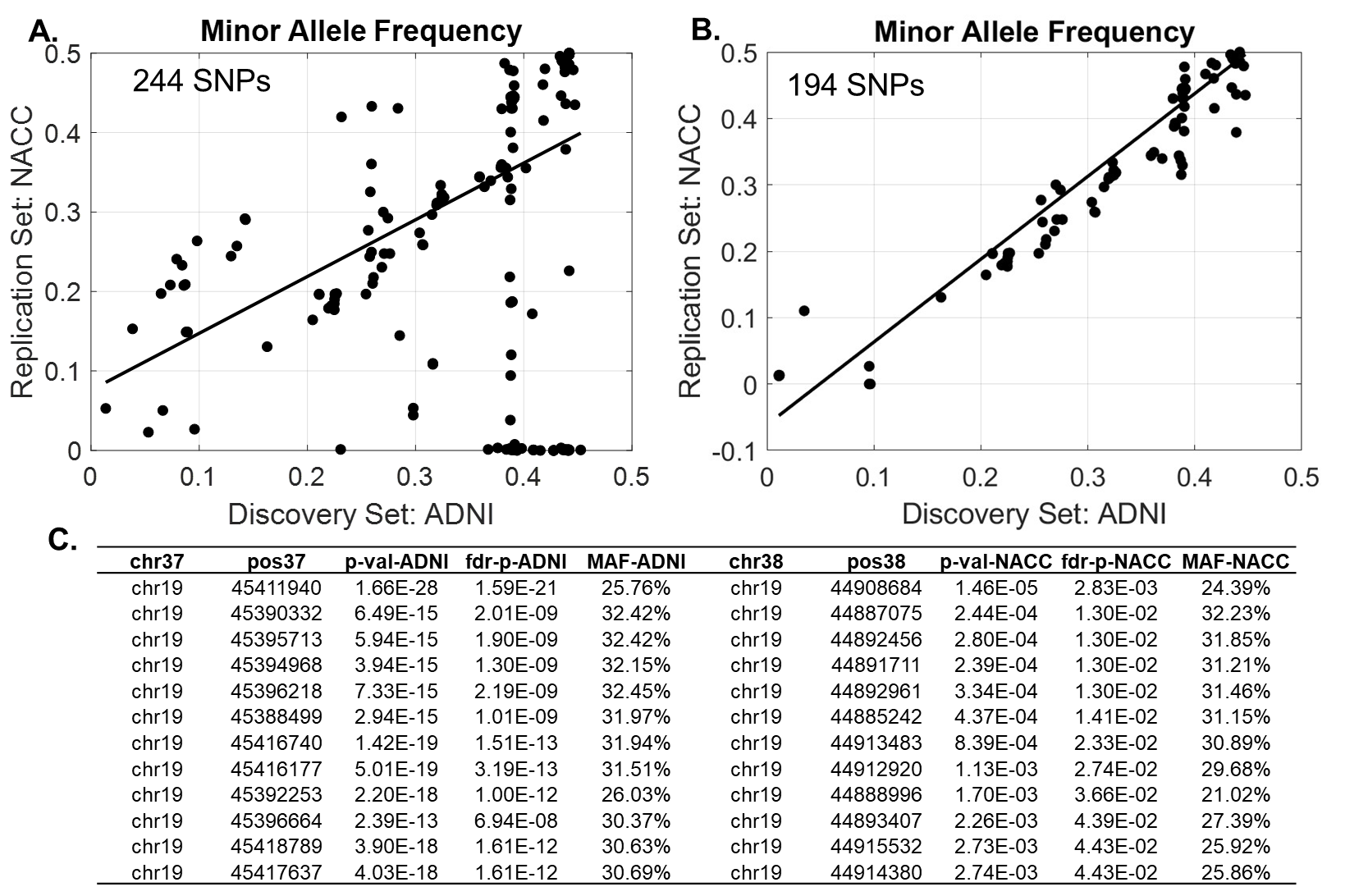

Supplement Fig. 1. Replication analyses of 244 SNPs identified using ADNI participants on the binary phenotype of clinical impairment status. (A) Minor allele frequencies of these 244 SNPs between ADNI (x-axis) and NACC (y-axis) participants. (B). Minor allele frequencies of 194 SNPs with a consistent MAFs in both databases. (C). 12 SNPs that retained to be significant at *p_FDR_*<0.05, corrected over 194 SNPs.

### Supplement Material 2. Longitudinal GWAS using only amyloid positive subjects.

We further limited our analyses to use longitudinal visits from amyloid positive participants on the phenotype of clinical impairment status in AD. Participants’ amyloid positivity status were determined from their latest PET amyloid scans. Participants with a whole-brain SUVR normalized to cerebellum greater than 1.11 were classified as amyloid positive participants. A total number of 4,052 clinical visits from 620 participants were included in this analysis.

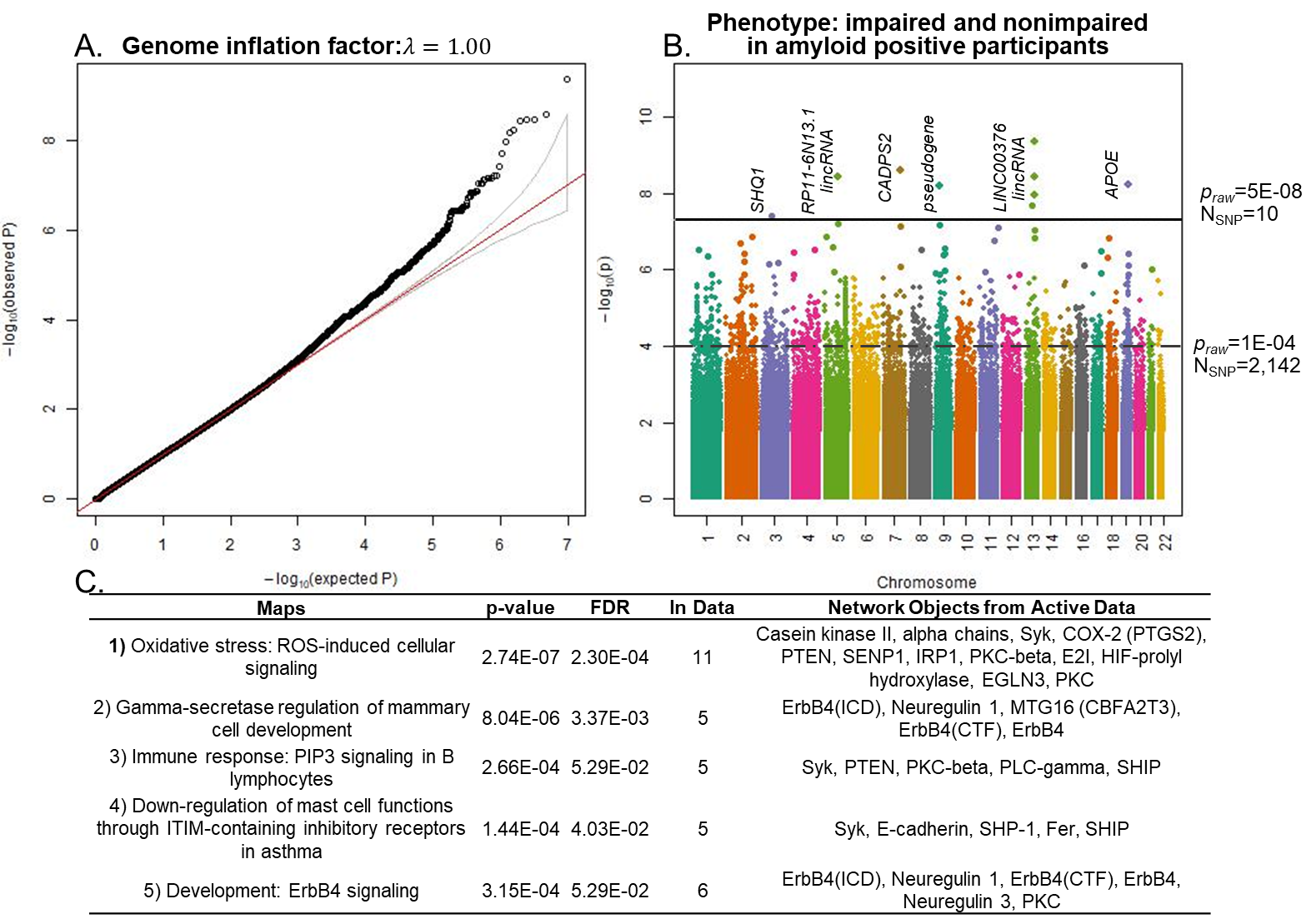

As shown in Supplement Fig. 2 above, ten SNPs reached genome-wide significance level of *p_raw_*<5E-08 and 2,142 SNPs reached a significance level of *p_raw_*<1E-04 (B). Pathway analyses on these 2,142 SNPs suggested that oxidative stress to be the most disrupted biological pathway associated with these variants (C).
